# Soil organic carbon fractionation and metagenomics pipeline to link carbon content and stability with microbial composition – First results investigating fungal endophytes

**DOI:** 10.1101/2021.12.19.473394

**Authors:** Wolfram Buss, Raghvendra Sharma, Scott Ferguson, Justin Borevitz

**Affiliations:** Research School of Biology, Australian National University, 134 Linnaeus Way, 2601 Canberra, Australia

**Keywords:** Metagenome, carbon fractionation, fungal inoculant, carbon sequestration, negative emission technology

## Abstract

Society needs to capture gigatons of carbon dioxide from the atmosphere annually and then store it long-term to limit and ultimately reverse the effects of climate change. Bringing lost carbon back into agricultural soils should be a priority as it brings the added benefit of improving soil properties. Linking soil organic carbon (SOC) fractions of different stability with soil microbial composition can help understand and subsequently manage SOC storage. Here we develop a pipeline for evaluating the effects of microbial management on SOC content using rapid and low-cost SOC fractionation and metagenomics approaches. We tested the methods in a wheat pot trial inoculated with 17 individual endophytic fungal isolates. Two fungi increased total SOC in the area under the plant stem by ~15%. The fractionation assay showed that the medium stability soil aggregate carbon fraction (AggC) was increased by one of these fungi (+21%) and the chemically recalcitrant proportion (bleach oxidation) of AggC by the other (+35%). Both fungi increased mineral-associated organic carbon (MAOC), the long-term SOC storage, by ~10%. We used rapid, portable, low-cost, whole metagenome long read sequencing to detect a shift in the microbial composition for one of the fungi-inoculated treatments. This treatment showed a more diverse microbial community and a higher quantity of DNA in soil. The results emphasise the link between composition and abundance of soil microorganisms with soil carbon formation. Our dual carbon fractional and metagenomic analysis pipeline can be used to further test the effects of microbial management and ultimately to model the soil factors that influence SOC storage, such as nutrient and water availability, starting SOC content, soil texture and aggregation.

## 1 Introduction

Soil organic carbon (SOC) sequestration is crucial for reducing atmospheric carbon dioxide levels and stabilising the climate (Paustian et al., 2016; Smith, 2016). SOC has a positive environmental and agricultural impact. It reduces nutrient leaching and increases water retention (Smith, 2016); SOC levels are positively correlated with crop yields globally (Oldfield et al., 2019). To benefit from these effects long-term and to influence atmospheric carbon dioxide levels on a societal relevant scale, SOC needs to be stored in a stable form to avoid re-release of the carbon on short-term.

There is good evidence that SOC persistence increases with higher soil aggregation (Peng et al., 2017; Rovira and Greacen, 1956; Six and Paustian, 2014) and that mineral-associated organic carbon (MAOC) is the long-term SOC reservoir (Hemingway et al., 2019). Most researchers also agree that the least stable carbon pool is particulate organic carbon (POC), which is derived from plant litter and roots that are in their early stages of decomposition (Abramoff et al., 2018; Cotrufo et al., 2019; Lavallee et al., 2020; Poeplau et al., 2018). Newer SOC process models contain measurable carbon pools where these three main SOC fractions are typically distinguished: POC, carbon protected in aggregates (AggC) and MAOC (Abramoff et al., 2018; Robertson et al., 2019).

Spatial SOC variability in the field can be as high as 4-fold (Robertson et al., 1997) and can decrease substantially with depth. Background SOC stocks in soils are 30-90 t C ha^−1^, while changes to agricultural practice typically increase the SOC stock by <0.5-1 t C ha^−1^ yr^−1^ (Paustian et al., 2019). Therefore, it is challenging to detect statistically significant changes in SOC in the short term (within a year) in the field (Paustian et al., 2019). Consequently, large sets of soil samples need to be analysed to increase the likelihood of detecting statistically significant differences in SOC content. SOC fractionation assays can help separate signal from noise by removing the labile, POC fraction that both sharply increases, e.g., when crop residues are returned to the field, and also quickly decreases due to microbial decomposition. Furthermore, separating AggC from MAOC can reveal alternative sequestration mechanisms.

Plant carbon delivered below ground through rhizodeposits is converted into SOC much more efficiently than from plant litter that decomposes on the soil surface (Sokol and Bradford, 2019). However, a large proportion of gross rhizodeposition, on average ~55%, is decomposed and released back as CO_2_ (Pausch and Kuzyakov, 2018). Microorganisms are responsible for the respiration of plant carbon but they also convert carbon into more stable SOC forms (Liang et al., 2019). Therefore, increasing the retention of plant rhizodeposits in soil in the form of AggC and MAOC through management of microbial communities in the plant rhizosphere could be a way to increase carbon sequestration in agricultural systems (Kallenbach et al., 2019), while also prodiving benefits on nutrient cycling and water retention.

The addition of endophytic fungal isolates for increasing SOC content has shown some success (Alves et al., 2021; Mukasa Mugerwa and McGee, 2017). However, the mechanisms responsible for this change in SOC content and how soil microbial composition and stable SOC formation are linked are mostly unknow. Many factors can potentially influence the plant-fungi-soil interplay and hence SOC accumulation, such as soil nutrient and water availability, soil texture, mineral composition or starting SOC content. Therefore, to enable predictable effect of microbial management, high-throughput and cost-effective assessments of soil biology and chemistry are required, most notably unbiased microbiome analyses that determine the whole suite of soil organisms in soil. These analyses should subsequently be linked to understand the microbial composition that drives stable SOC formation.

The objectives of this study were to (i) develop a pipeline for rapid screening of SOC stability and microbial composition, (ii) assess the effect of endophytic fungal isolates in wheat on SOC content and (iii) investigate covariation of SOC shifts with changes in the microbiome.

## 2 Materials and Methods

### 2.1 Pot trial

The soil used in the trial was a loam from central western New South Wales in Australia with a starting SOC content of 1.1%, pH of 7.34 and a total N content of 0.20%. Details of the soil can be found in SI Table 1.

The soil was filled into 11 cm diameter and 11 cm high, round pots. The endophytic fungi were grown on potato dextrose agar plates and small plugs from the plate were cut out and placed in a slight depression in the soil. Wheat seedlings (Condo variety) were pre-germinated and transplanted over the fungal plug. Overall, 17 fungal isolates were tested supplied by SoilCQuest (Orange, New South Wales, Australia). Of the fungi inoculants 14 were of the phylum Ascomycota, and one of the phyla Mucoromycota, Basidomycota and Zygomycota, respectively (SI Table 2). A randomised block design was used for eight replications per treatment including a control without fungal inoculation. The temperature and light profiles were adjusted following the actual conditions of the winter growing season of 2018 (May-November) in Parkes, New South Wales, Australia with a temperature profile of ~8-10ׄ°C at night and ~18-25°C during the day. The pots were watered regularly (~every 3 days) using tap water. At the end of the experiment, the wheat plants were cut above the soil level, dried and weighed.

### 2.2 Soil sampling and total carbon analysis

Soil cores were taken with 50 mL centrifuge tubes underneath the plant stem to a depth of ~3 cm. The soil was dried, followed by gentle tapping with a pestle in a mortar and sieved to < 2 mm to remove gravel and roots. This soil sample from below the stem is referred to as the ‘inner-rhizosphere soil’ as it contains the main wheat roots. A subsample was ground, and the total carbon content was analysed with a VarioMax 3000 (Elementar, Germany). From the treatments with significant increases in total carbon levels in the ‘inner-rhizosphere’, including the control, three sub samples were taken to assess the change in total SOC in the whole pot (here termed ‘outer-rhizosphere’ soil). The locations within the pot were selected randomly, excluding the central area where a sample was already taken for the ‘inner-rhizosphere’ soil. Figure 1 shows a schematic of the soil sampling regime.

**Figure 1:**
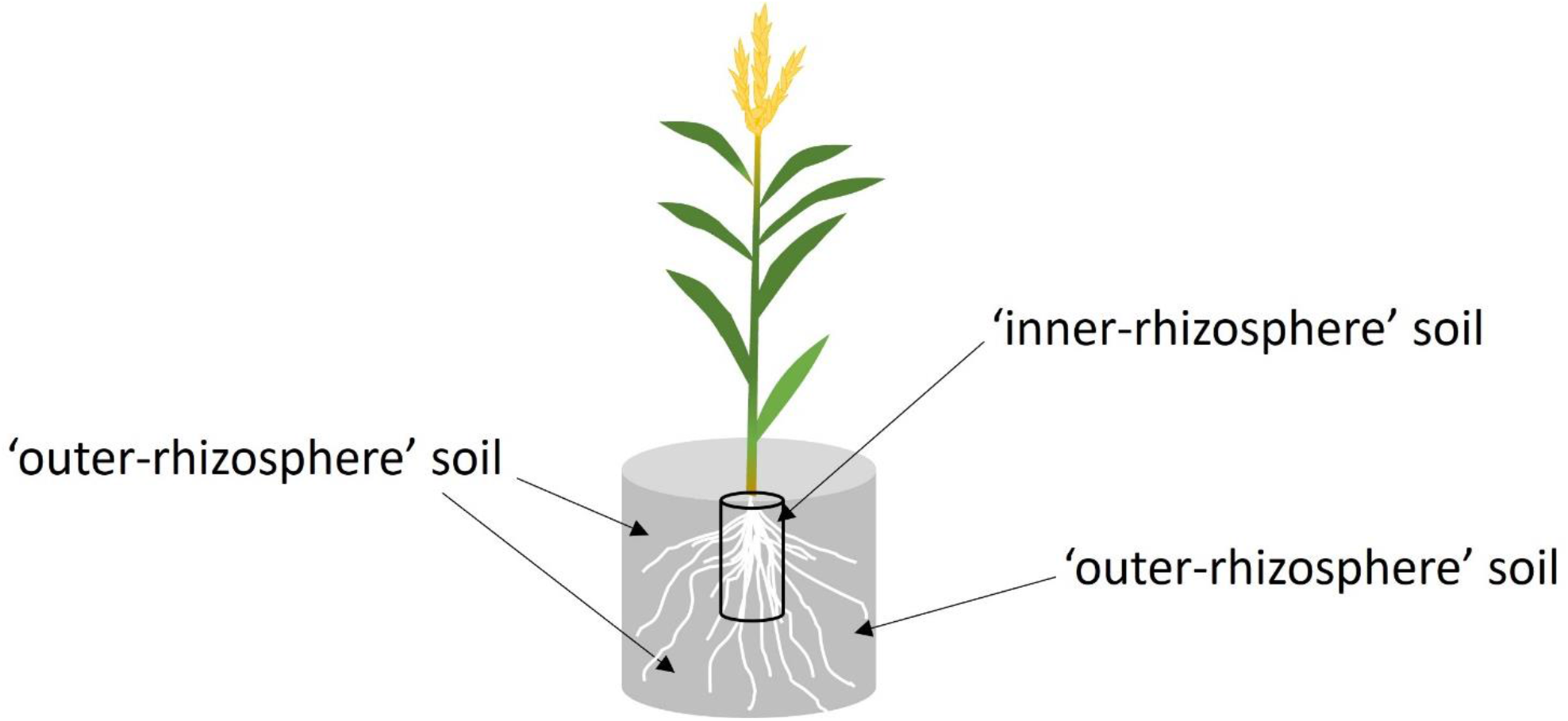
Soil sampling regime in wheat pot trial. One ‘inner-rhizosphere’ soil sample was taken in all pots for total C analysis. In treatments with a significant increase in total C content in the ‘inner-rhizosphere’ soil, three additional soil samples were taken from the ‘outer-rhizosphere’ area.

### 2.3 Soil fractionation

#### 2.3.1 Development and rationale

SOC fractionations separate the soil into carbon pools of different stability. Many different fractionation techniques that are described in the literature (Poeplau et al., 2018). We decided to use carbon pools that could be used for subsequent modelling of SOC dynamics over time and space to predict changes in management practices. Yet we aimed to develop a cost-effective and rapid approach to enable analyses of large number of samples in follow-up experiments, so various adjustments to existing methods were performed allowing for higher throughput analysis.

The soil fractionation assay was developed based on Poeplau et al. who compared and evaluated 20 different studies and concluded that dispersion of aggregates followed by size fractionation and a subsequent density separation yielded the best results (Poeplau et al., 2018). We also added a step to determine chemical recalcitrance, as suggested in the study.

Some fractionation protocols start with a density separation, followed by full disaggregation and a size fractionation to separate out ‘unprotected’ POC (Poeplau et al., 2018) We chose to first perform wet-sieving to avoid contamination of the aqueous fraction (DOC) with the high concentration of salt used for density separation. This enables sub-samples of the aqueous fraction to be analysed for pH, electric conductivity (EC) and water-extractable nutrients. A soil-to-water ratio of 1:5 was selected for the initial disaggregation as this is the ratio typically used to determine pH and EC in soils. Weighing out soils and performing extractions are labour and time intensive. This one-step procedure allows the analysis of multiple parameters on the same soil sample that can be correlated with each other reducing challenges with soil heterogeneity. This data can subsequently be the input for statistical models to link soil chemical parameter, soil carbon and microbial composition.

In short: after initial, gentle disaggregation of the soil sample by shaking with water, we wet-sieved the sample to separate coarse from fine material. Subsequent density separation of the coarser material spilt the sample into AggC (heavy) and POC (light). Finally, the fine fraction was separated via another size/density separation into MAOC (heavy) and dissolved organic carbon (“light”; dissolved in water; not measured). A flow-chart is presented in Figure 2.

**Figure 2:**
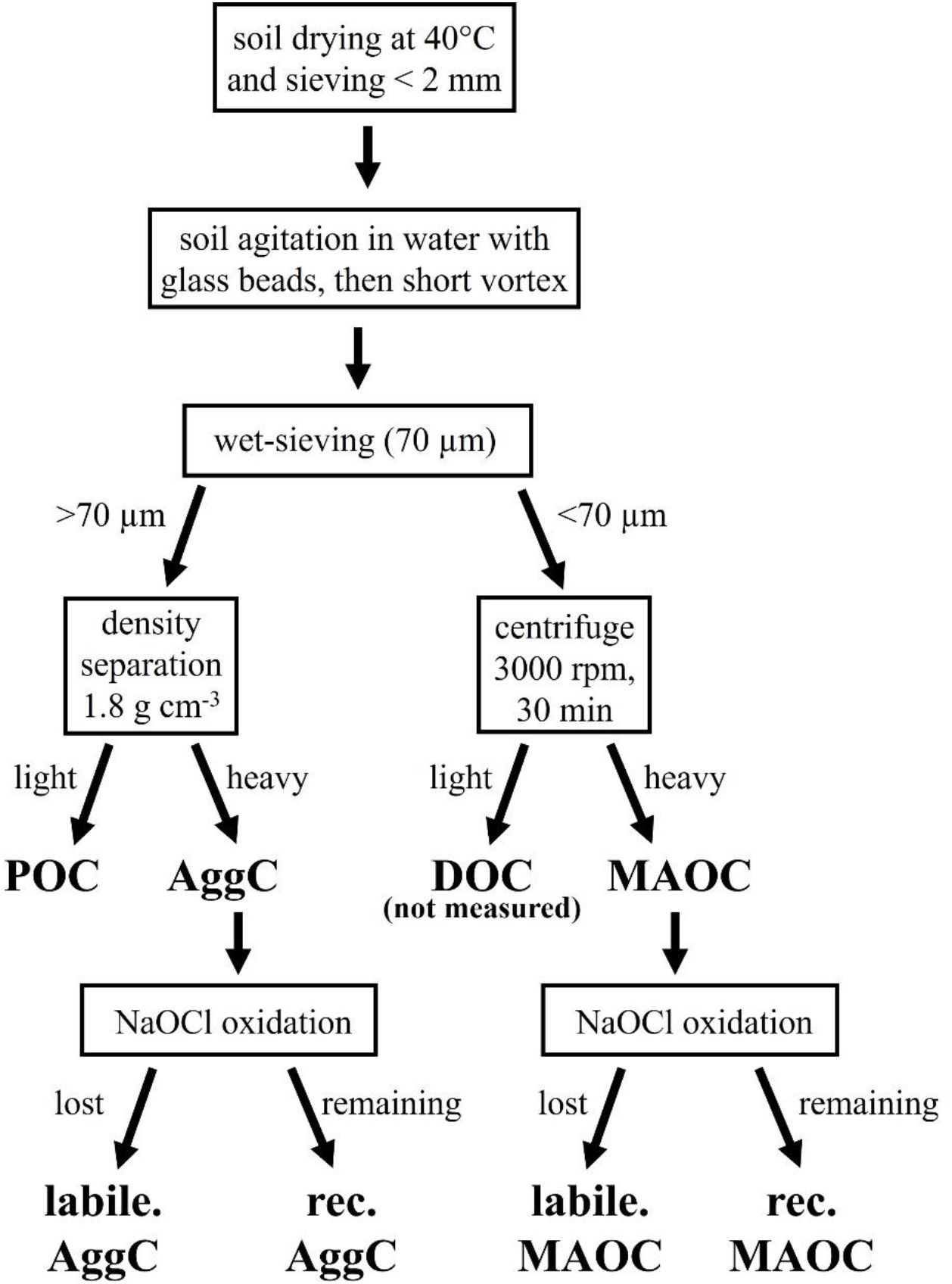
Schematic of soil carbon fractionation protocol developed and used in this study.

#### 2.3.2 Protocol

Samples of the treatment with a statistically significant increase in total carbon levels in the ‘inner-rhizosphere’ were fractionated without prior soil grinding.

The following steps were performed: (i) 10 g of soil was shaken with 25 mL deionised water and 2 glass beads for 5 min at 80 rpm in 50 mL falcon tubes on an orbital shaker with the falcon tubes lying horizontally; (ii) wet sieving was performed using molecular sieves of 70 μm size that fit onto 50 mL falcon tubes, which accelerated the sieving process from >20 min per samples to <5 min (a larger cut-off than the typical 50, 53 or 63 μm was used since less water than for comparable wet-sieving methods was used and hence separation is less complete); (iii) the soil left on the sieve was washed back into the falcon tube with 12.5 mL of deionised water and the tube was vortexed for 2 s, then the sample was sieved again and the process was repeated, which resulted in a total water-to-soil ratio of 5:1 (50 mL of water to 10 g of soil); (iv) for separating POC and AggC sodium iodide adjusted to a density of 1.8 g cm^−3^ was used as recommended in Sohi et al. (Sohi et al., 2001), who tested 1.6, 1.7 and 1.8 g cm^−3^, and which is also used in the SOC model by Robertson et al. (Robertson et al., 2019); (v) the samples were centrifuged to ensure the AggC fraction remains at the bottom of the tube; (vi) the POC and AggC fraction were separated by decanting the content of the tube after centrifugation onto a Whatman No. 2 filter paper through turning the tube by 360° while decanting; (vii) the filter paper was washed with deionised water, dried and weight to determine the POC fraction; (viii) the MAOC fraction was separated from the dissolved organic carbon fraction via centrifugation at 3000 rpm for 30 min (instead of using time-intensive vacuum filtration to <0.45 um) as suggested before (Robertson et al., 2019); (ix) to remove sodium iodide residues, the AggC fraction was washed with deionised water, though only once to avoid carbon loss (Plaza et al., 2019).

The POC, AggC and MAOC fractions were dried, weighed and the total carbon contents were analysed. The data were expressed as carbon in weight percent that is associated with the respective SOC fraction.

#### 2.3.3 Rapid recalcitrance test

In addition to the fractionation assay, which assesses the physical protection of SOC from degradation, we also conducted a chemical recalcitrance test to “mimic strong enzymatic decay and isolate the oxidation-resistant SOC fraction” as outlined in Poeplau et al. (Poeplau et al., 2017).

Our method was adapted from Poeplau et al. (Poeplau et al., 2017) using 6% sodium hypochlorite (NaOCl) that was added to 1 g of the AggC and MAOC fractions. The samples were incubated in the lab for 20 hours and subsequently centrifuged. The NaOCl solution was decanted, and the treated soil was washed once with deionised water, followed by centrifugation and decanting. Finally, the samples were weighed, and the remaining carbon content was measured.

### 2.5 Soil metagenomics

The soil metagenomics approach was based on existing techniques that were adapted to allow for rapid, in-house tested for low-cost. The assay can be performed for <US$ 20 per sample depending on sequencing depth. A sub sample (350-400 ug soil) was taken from the ‘inner-rhizosphere’ soil samples and subjected to DNA extraction using the DNeasy PowerSoil Pro Kit (Qiagen, Hilden, Germany) using the manufacturer’s instructions. The DNA concentration in each sample was measured using a ND-1000 spectrophotometer (NanoDrop). Subsequently, the DNA was barcoded using the Oxford Nanopore rapid barcoding kit following manufacturer’s instructions using 1 μl of rapid barcode and eluting with water. The 24 samples were pooled in 2 groups of 12 samples and each run on a flongle (Oxford Nanopore Technology, Oxford, UK).

Data obtained from the flongle sequencer was converted into DNA sequences (basecalled) and demultiplexed with Guppy (version: 5.0.7). Sequence libraries were next curated, removing all short (<50 bp) and low quality (<q7) sequences using NanoPack (De Coster et al., 2018). Curated sequence libraries were blasted against the NCBI nucleotide database version 5 (downloaded on 2021-09-30) (Sayers et al., 2021). Finally, the taxonomic ID for the single best blast hit per sequence was extracted and using the python3 library The Environment for Tree Exploration (Huerta-Cepas, 2016) the full taxonomic classification (phylum, class, order, family, genus, and species) of each sequence match was obtained. No match sequences and sequences that were assigned to species that were only detected once in a sample were filtered out. Sequences matching a phylum observed in only a single sample were also filtered out. The common phylum abundance in each sample was normalized to the total filtered sequences in each sample. The data were used to perform a principal component analysis (PCA) in R (prcomp; scaled and centred) and the output was subsequently visualised (autoplot; package ggfortify).

## 3 Results

### 3.1 Changes in total SOC due to fungal endophytes

The carbon (C) content in the area underneath the stem of the plant where the bulk of the plant roots are located (here termed ‘inner-rhizosphere’) increased significantly by 15% and 17% in isolates 112 and 1852 compared to the control, respectively (ANOVA with Dunnett’s test; Figure 3). Changes with other fungal treatments were not significantly different among biological pot replicates and were not further analysed. Subsequently, we analysed the C content in three additional locations that were randomly picked within the pot in isolates 112, 1852 and the control. These ‘outer-rhizosphere’ samples did not reveal a significant increase in carbon content in isolate 1852 but an increase in isolate 112 by 8% (p = 0.036) (Figure 4).

**Figure 3:**
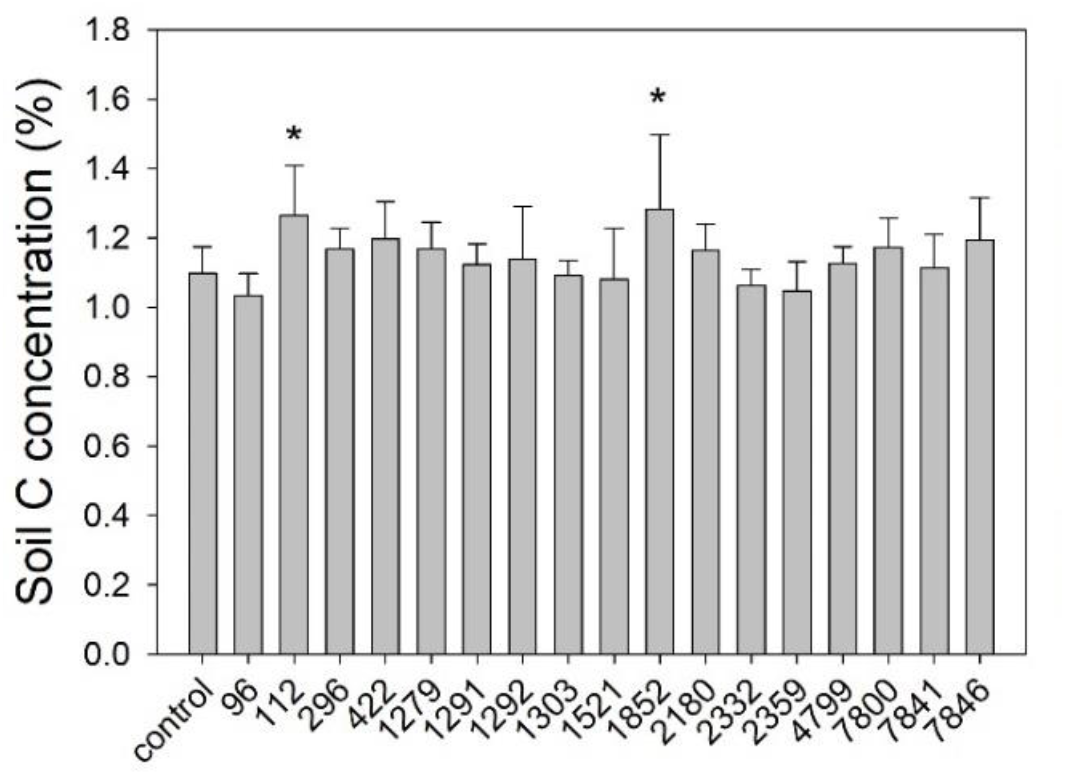
Total carbon content in soil samples taken from the area underneath the plant stem at harvest (‘inner-rhizosphere’). Mean and standard deviation (SD) of eight replicates reported. *statistically significant differences determined by one-way ANOVA followed by Dunnett’s post hoc test comparing fungi treatments to controls (p < 0.05).

**Figure 4:**
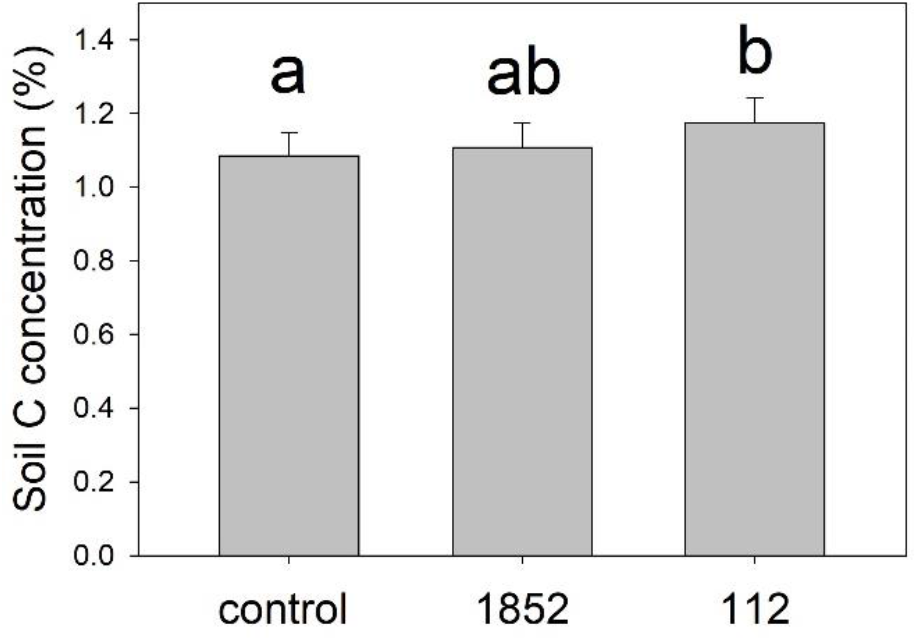
Total carbon content in the ‘outer-rhizosphere’ soil in control and two isolate treatments. Three soil samples were taken from each of the eight pot replicates randomly as outlined in Figure 1, and each analysed for total C. The mean of the three samples per pot was calculated and subsequently mean and standard deviation (SD) of eight pot replicates reported. An ANOVA followed by Tukey post-hoc test was performed.

### 3.2 Soil fractionation assay to detect carbon of different stability

The ‘inner-rhizosphere’ soils of the control treatment and fungal isolates 112 and 1852 were subsequently fractionated into POC, AggC and MAOC. The N associated with the POC fraction increased by 93% on average in the samples treated with fungi 112 (Figure 5B), while the C content in the POC fraction was not different to the control (Figure 5A). This decreased the C/N ratio in the POC fraction significantly by 48% (Figure 5C). Among the three soil fractions, the POC fraction has the highest C/N content by far (control C/N ratio 52) as also reported in other studies (Blume et al., 2016).

**Figure 5:**
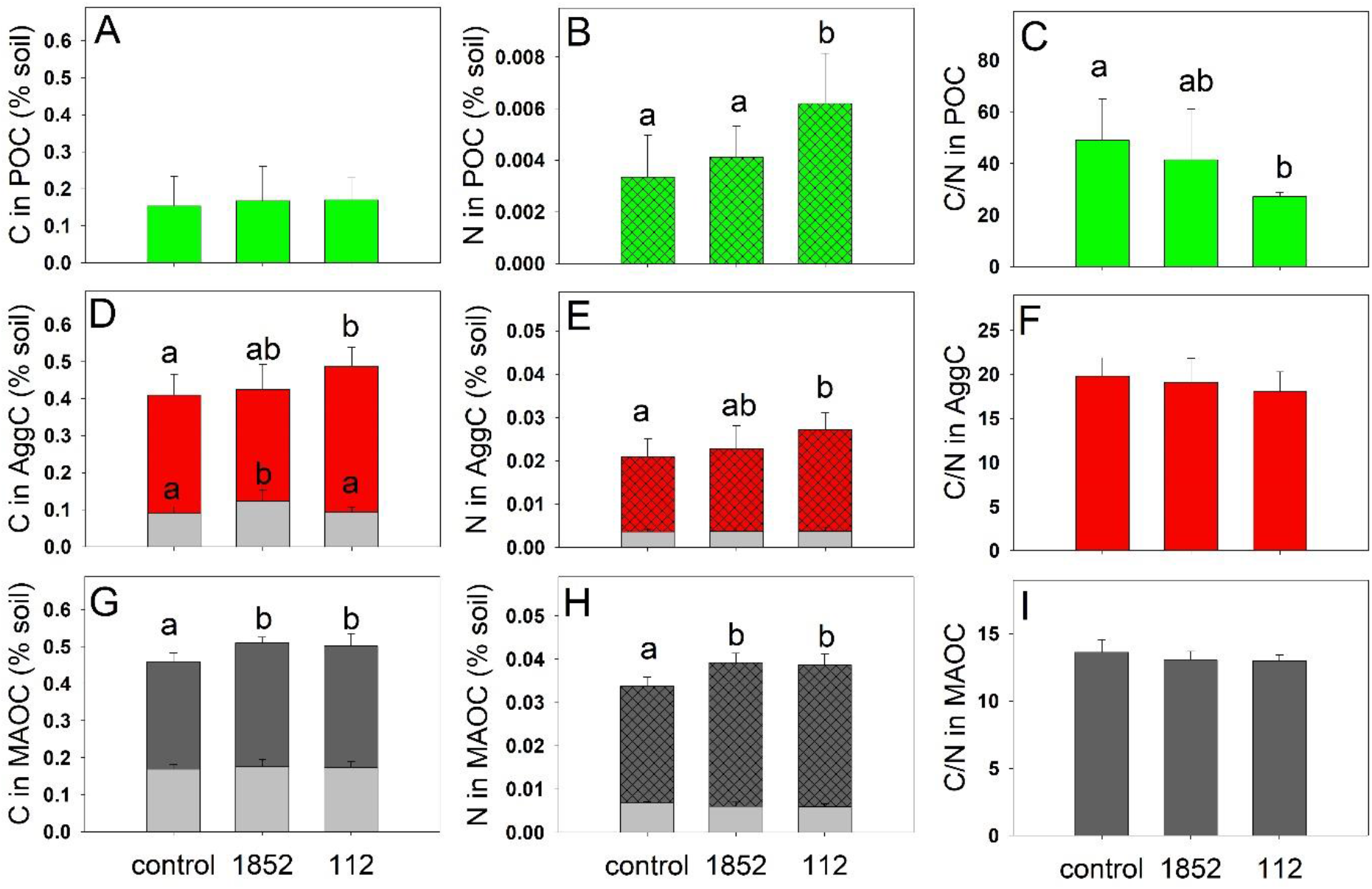
Contents of carbon (A, D, G) and nitrogen (B, E, H), C/N ratios (C, F, I) and carbon and nitrogen recalcitrance (grey bars in D, E, G and H) in the ‘inner-rhizosphere’ soil that was separated into three soil carbon fractions. Three treatments were investigated: control and two fungal isolates. (A, B, C) Particulate organic carbon (POC), (D, E, F) aggregate carbon (AggC) and (G, H, I) mineral-associated organic carbon (MAOC). Chemical recalcitrance (bleach oxidation test) of carbon and nitrogen within AggC and MAOC fractions are highlighted by grey bars. Soil samples were taken from the area underneath the plant at harvest (‘inner-rhizosphere’). Mean and standard deviation (SD) of eight replicates reported. Different letters indicate significant differences among the treatments using one-way ANOVA followed by Tukey post-hoc test.

The carbon content associated with the AggC fraction increased by 21% in the 112-isolate treatment (Figure 5D) and N increased by 34% (Figure 5E). Both isolates significantly increased the C and N associated with the MAOM fraction by 8-10% (Figure 5G) and 14-16% (Figure 5H), respectively. The C/N ratios did not change significantly with fungal inoculation in the AggC and MAOC fractions (Figure 5F and I).

The C and N contents associated with the AggC and MAOC fractions correlated well with each other (Figure 6B, C) confirming a constant ratio of C and N in all samples. In the POC fraction, the C and N contents correlated well in the samples of treatments 112 and 1852 (except for one outlier in 1852) (Figure 6A). However, the C and N contents in the control samples did not follow the same relationship (Figure 6A) confirming the results from the average C/N ratio in these treatments.

**Figure 6:**
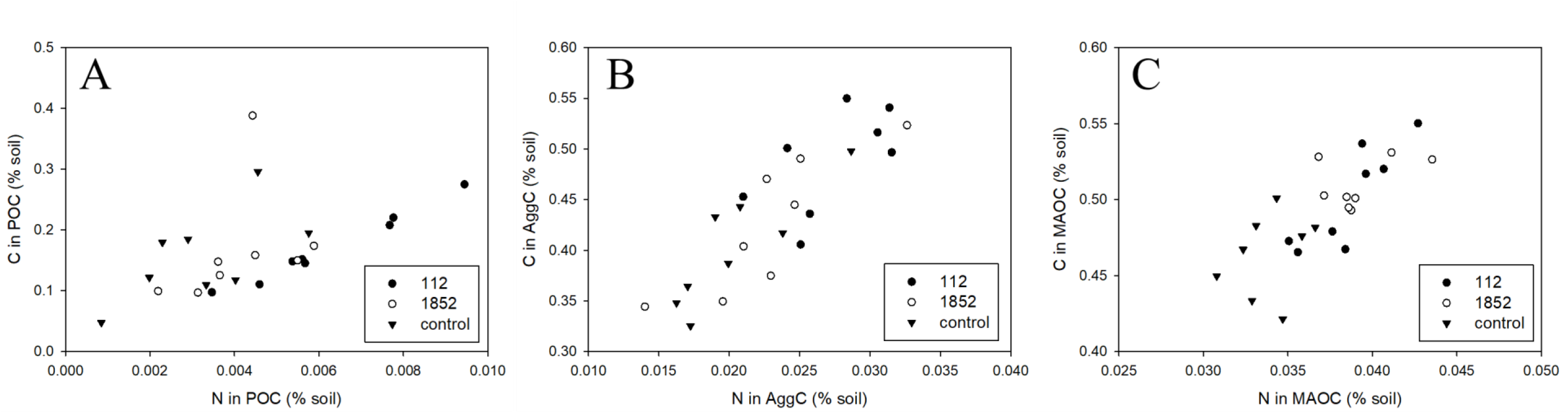
Contents of carbon and nitrogen associated with (A) particulate organic carbon (POC), (B) aggregate carbon (AggC) and (C) mineral-associated organic carbon (MAOC) in control and two fungal isolate treatments. Soil samples were taken from the area underneath the plant at harvest (‘inner-rhizosphere’).

We tested the AggC and MAOC fractions for chemical recalcitrance via a bleach oxidation assay and found a significant increase in recalcitrant C in the AggC fraction in soil treated with isolate 1852 by 35% but no increase in 112-treatments (Figure 5D; grey bars). No significant change in recalcitrant C was detected within the MAOC fraction in either of the fungi-treatments.

### 3.3 Shift in soil microbial composition detected by whole metagenome sequencing

DNA was extracted from 0.4 g of inner-rhizosphere soil for eight replicates of each control and fungal treatments. The DNA concentrations in the soil samples of the control and treatments 1852 and 112 were 29.6 ± 7.4, 35.1 ± 9.5 and 40.5 ± 5.8, respectively. The DNA concentration in treatment 112 was significantly higher than in the control (p = 0.004; 2-sided, equal variance t-test) suggesting a higher abundance of DNA in the 112-inoculated soil sample.

We performed a PCA on phylum level microbial composition using DNA extracted from each individual pot that was blasted against the NCBI nucleotide database. The PCA plot demonstrates a shift in composition due to the addition of fungi 112 (Figure 7A). The soil samples of several pots that were inoculated with 112 separate from the remaining samples through a shift on principal components (PC) 1 and 2 (Figure 7A). This shift is in parts driven by a higher abundance of the phylum Ascomycota, which is the phylum of both added fungi. This phylum was detected in six of the eight replicates of the 112-treatment, averaging ~0.5% of the sequences detected in the soil, however, it was not detected in any samples of the 1852 treatments (or the control) (Figure 8B). The average microbial composition of the common phyla in treatment 112 only changed marginally (Figure 8A) but the rarer phyla comprise a higher percentage of the total microbial composition in samples treated with fungi 112 (Figure 8B). The results indicate that the growth of fungi 112 in the rhizosphere stimulated a more diverse microbial composition.

**Figure 7:**
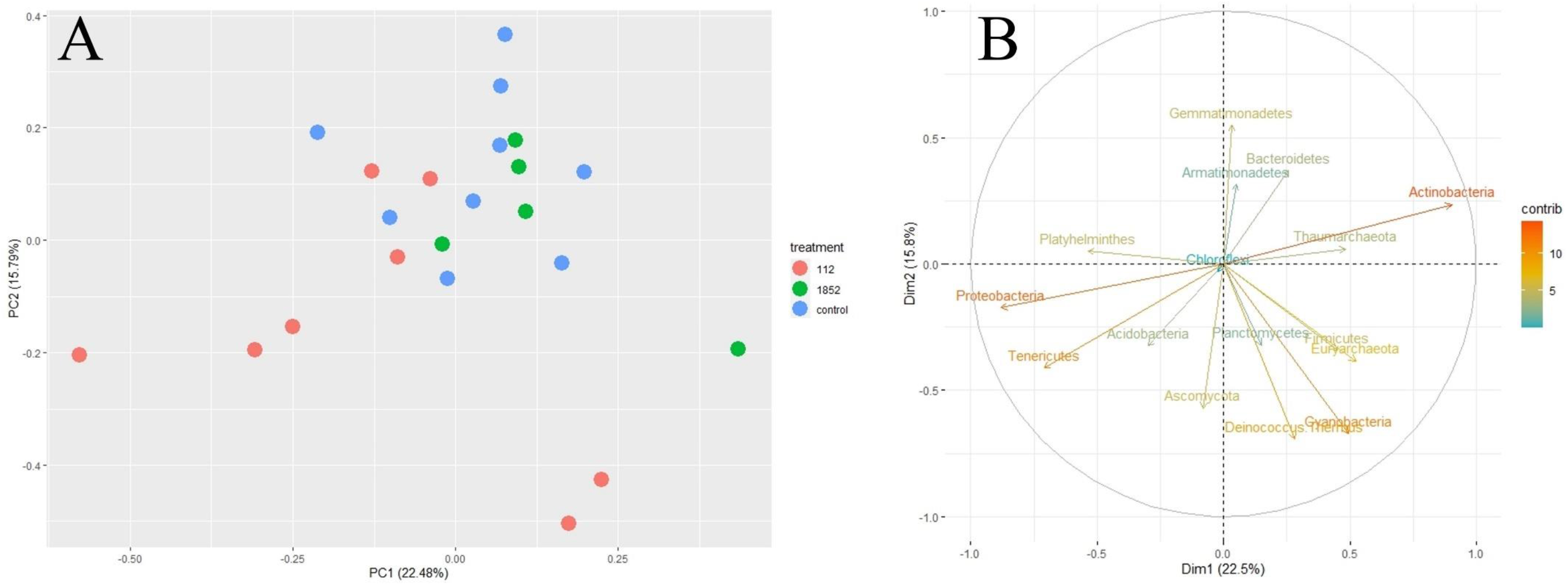
Principal component analysis (PCA) of soil microorganism composition as relative percentage of detected phyla in each pot of the plant growth experiment inoculated with fungi 112, 1852 or uninoculated (control). (A) Graph of individual samples and (B) graph of variables. DNA was extracted from soil samples taken from the area underneath the plant at harvest (‘inner-rhizosphere’).

**Figure 8:**
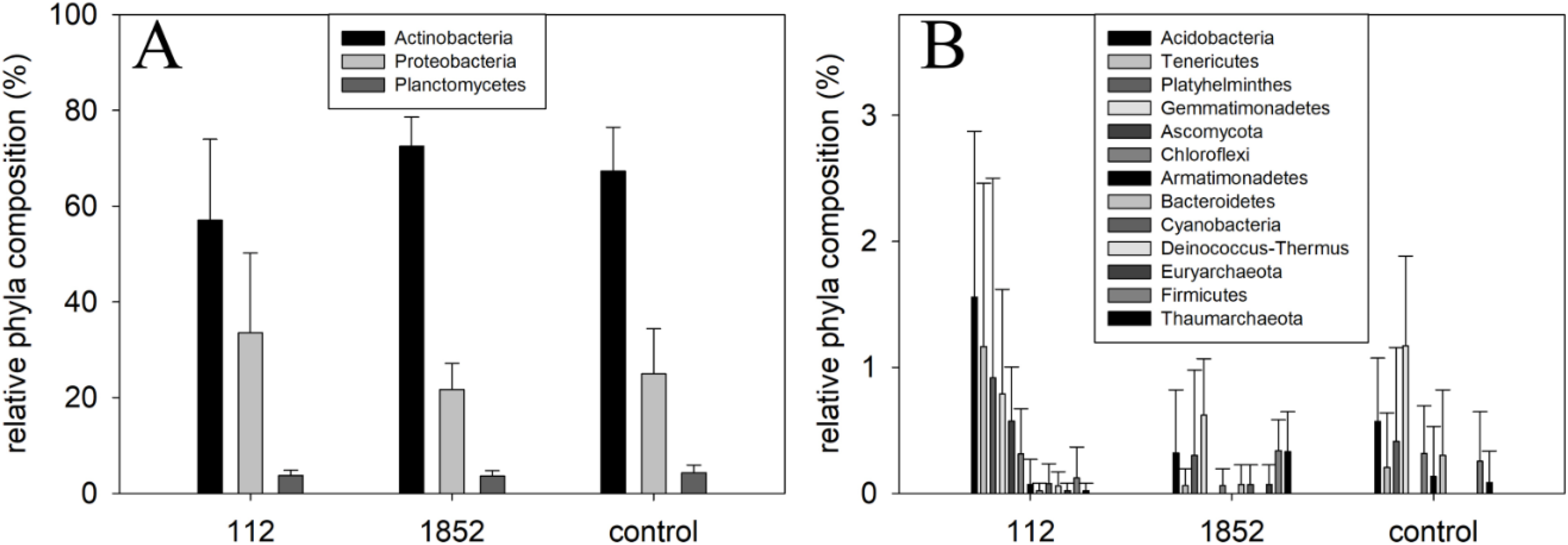
Soil microbial composition in the soil samples from the plant growth experiment inoculated with fungi 112, 1852 or uninoculated (control). (A) Common phyla and (B) rarer phyla (<3% of total composition). DNA was extracted from soil samples taken from the area underneath the plant at harvest (‘inner-rhizosphere’).

Isolate 1852, however did not separate from the control in the PCA plot. In contrast to soil samples treated with isolate 112, the samples with 1852 did not demonstrated an increase in total C in the ‘outer-rhizosphere’ and no increase in AggC (Figure 4D). This demonstrates that the metagenomics and SOC fractionation assay results align; in both cases the effect of 112 was more pronounced than the effect of 1852.

## 4 Discussion

In this study we detected an increase in SOC content after inoculation of wheat roots with fungal endophytes. The results demonstrate that even in the (small) pots used in this trial the effect on SOC content is spatially restricted decreasing with distance from the primary plant roots where most of the plant exudation occurs. For detecting differences in SOC stocks due to fungal inoculation analysing the ‘inner-rhizosphere’ is an effective way of detecting small differences in short-term experiments. Yet, this does not accurately represent the SOC content in the whole pot and because the effect is spatially explicit, neither measurement should be used to extrapolate the effect across an agricultural field. Actual field measurements are needed to establish temporal and spatial SOC dynamics.

There are various explanations for the SOC increase, i.e., increased plant carbon exudation, higher persistence of fungal carbon in soil and increased conversion efficiency of plant exudation into fungal biomass.

There is good evidence that plant photosynthesis is not source but sink limited (Gavito et al., 2019; Laheurte et al., 1990) and plant-associated microorganisms constitute a strong sink for plant exudates (Canarini, 2019; Gavito et al., 2019). Our results indicate a higher abundance of endophytic fungi 112 in our experiment as shown by higher levels of DNA in soil and a shift in microbial composition towards the phylum of the added fungi. So, with the deployment of the fungal endophyte that takes carbon from the plant, photosynthesis increases without reduced plant growth (Ahkami et al., 2017; Gavito et al., 2019) (the plant biomass did not change with 112-inoculation; SI Figure 1). Subsequently extra carbon is provided to the soil in the form of plant exudates.

This higher level of plant exudation could directly affect the SOC level through accumulation in soil or the plant exudates are first converted into fungal biomass/necromass through anabolism (Sokol et al., 2019; Sokol and Bradford, 2019). The additional C and N (extra relative to the control) associated with AggC and MAOC in soils treated with isolates 1852 and 112 had a C/N ratio of 9.4 and 8.5 and 12.4 and 7.7, respectively. On average, the C/N ratio of exudates in grasses is ~23 (Cardenas et al., 2021), while fungi and bacteria typically have C/N ratios of 10 and 4, on average (Liang et al., 2019), which could be evidence that the additional C and N is mostly fungal- and not plant-derived. The drop in C/N ratio in the POC fraction is further evidence for compounds with a low C/N ratio, such as fungi biomass.

This extra fungal biomass is likely more persistent in soil than plant-derived carbon, through more efficient sorption to minerals, through the formation of aggregates and through chemical recalcitrance (higher resistance against decomposition).

The fungal isolate 1852 increased the chemical recalcitrance of carbon in the AggC fraction. It is a dark pigmented fungus that likely has melanin present in the fungal cell wall that can remain in the soil for longer than necromass from non-melanised fungi (Fernandez et al., 2016; Fernandez and Kennedy, 2018; See et al., 2021). Melanin content in soil has been correlated with higher total soil C content (Siletti et al., 2017). Chemical recalcitrance is generally not longer considered a key mechanism for long-term SOC protection (Hemingway et al., 2019; Lehmann and Kleber, 2015). However, recalcitrant carbon in the form of fungal cell wall components, such as melanin and chitin, within aggregates that protect it from abiotic (oxidation) and biotic (microorganism) attacks could protect carbon over the medium-term (Cotrufo et al., 2013; Li et al., 2018). There is good evidence that microorganisms can self-assemble aggregates that occlude and hence protect their biomass and necromass (Rabbi et al., 2020; Throckmorton et al., 2015). This can explain the increase, or at least a proportion of the increase, in the AggC fraction in our study. This protection, however, would only be maintained in no-till systems where aggregates are not disturbed on an annual basis.

Microbial hotspots in small aggregates can also drive MAOC formation due to close proximity of microorganisms and mineral surfaces where necromass and microbial exudates can sorb to (Kravchenko et al., 2019). This sorption to mineral surfaces, as also observed in our study, can happen within hours (Buckeridge et al., 2020; Creamer et al., 2019) and there is evidence that microbial envelopes preferentially accumulate in the soil, sorbed to mineral surfaces (Kasanke et al., 2021; Liang et al., 2019; Miltner et al., 2012; Schurig et al., 2013).

The added fungi could also have a higher conversion efficiency of plant carbon (exudates) into fungal biomass than existing microbial communities, lowering plant-carbon losses (Anthony et al., 2020; Kallenbach et al., 2019, 2016). Proving this exudate-to-microbial biomass hypothesis would require mechanistic investigations that are outside the scope of this study but would be important future work.

All mechanisms described here likely play a role in increasing the SOC content: with increased growth of the fungal inoculant, more exudates are released by the plant roots; more plant exudation attracts a higher abundance and diversity of microorganisms (Baumert et al., 2018); this increased microbial activity subsequently creates more aggregates and protect the subsequent necromass from decomposition, increasing SOC content (Baumert et al., 2018); and mineral sorption protects microbial fragments on long-term.

More than half of SOC is comprised of microbial fragments and therefore, increasing the microbial biomass formation, microbial turnover and protection of necromass drive SOC accumulation (Liang et al., 2019). Microbial management in agricultural soils, microbial inoculation being one method, increasing plant exudate diversity another (Lange et al., 2015), could become a key strategy for climate change mitigation while improving agricultural productivity.

## 5 Conclusions

In this manuscript, we demonstrate the effectiveness of tools to rapidly and cost-effectively screen soil samples for SOC stability and microbial composition. Our low-cost metagenomics approach is unbiased to particular genetic loci and microbial species and was able to detect differences resulting from fungal inoculation in this study. The results from our SOC fraction assay demonstrates that certain fungal inoculations can increase the storage of plant carbon in a stable form. Analysing the aqueous portion of the soil fractionation assay for EC, pH and nutrients could be another addition to this assay. This would allow linking soil chemistry with soil biology and carbon of different stability. Considering the spatial and temporal variability of SOC content and its dependence on factors, such as soil nutrient status, the rapid screening tools described here are essential to process the large number of samples needed to calibrate models that are ultimately able to predict such changes.

## Acknowledgement

The funding for this project was provided by an ANU grand challenge grant with financial support and fungi inoculants from SoilCQuest2031.org (Orange, New South Wales, Australia). We acknowledge assistance from Austin Bird for lab and technical support.

