## Supplementary Information for "Soil organic carbon fractionation and metagenomics pipeline to link carbon content and stability with microbial composition – First results investigating fungal endophytes"

Wolfram Buss[a*]

Raghvendra Sharma[a]

Scott Ferguson[a]

Justin Borevitz[a]

[a] Research School of Biology, Australian National University, 134 Linnaeus Way, 2601 Canberra, Australia

SI Table 1: Properties of soil used in this study.

| parameter (unit) | value |
|---|---|
| total carbon (%) | 1.1 |
| total nitrogen (%) | 0.2 |
| Bray 1-Phosphorus (mg $kg^{-1}$ P) | 4.1 |
| pH (soil-to-water ratio 1:5) | 7.34 |
| electrical conductivity (dS $m^{-1}$) | 0.078 |
| calcium (%) | 67 |
| magnesium (%) | 24 |
| potassium (%) | 7 |
| basic texture | loam |
| basic colour | red |
| effective cation exchange capacity ($cmol_{+}$ $kg^{-1}$)) | 12 |

SI Table 2: Fungal isolates used in this study, supplied by SoilCQuest.

| fungal ID | phylum |
|---|---|
| 7846 | Ascomycota |
| 7841 | Ascomycota |
| 4799 | Ascomycota |
| 422 | Ascomycota |
| 1852 | Ascomycota |
| 7800 | Ascomycota |
| 2180 | Ascomycota |
| 1303 | Mucoromycota |
| 2359 | Ascomycota |
| 2332 | Ascomycota |
| 112 | Ascomycota |
| 96 | Ascomycota |
| 1292 | Ascomycota |
| 1291 | Ascomycota |
| 1521 | Basidiomycota |
| 1279 | Ascomycota |
| 296 | Zygomycota |

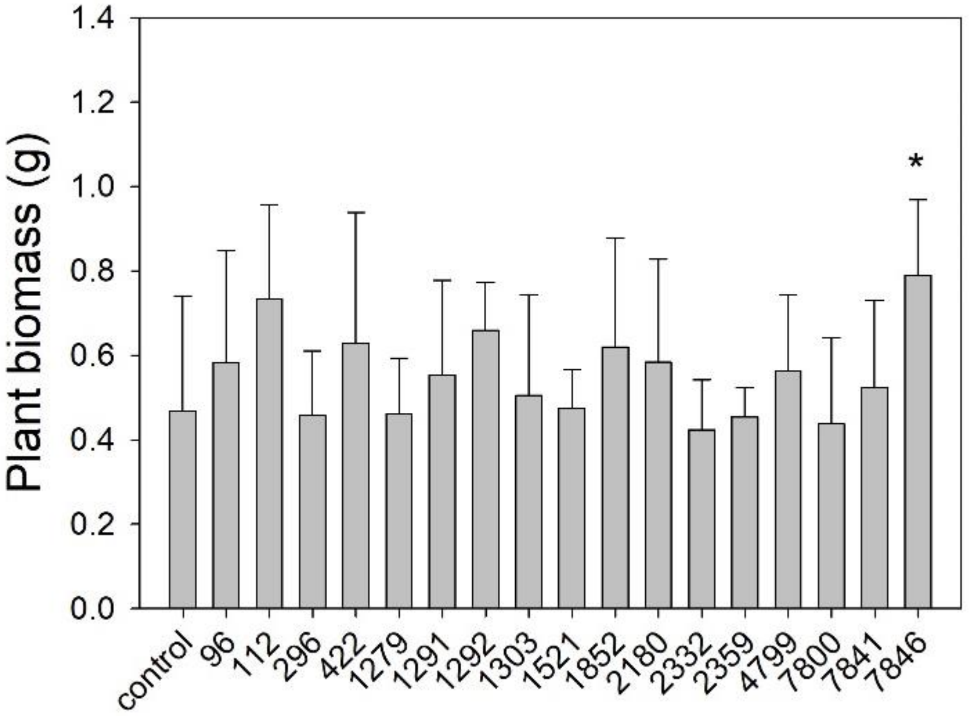


SI Figure 1: Plant biomass from the pot trial in this study.

SI Table 3: Total carbon content in soil cores under the plant stem (‘inner-rhizosphere’) and ‘outer-rhizosphere’ in three treatments (control and two fungal isolates) as mean and standard deviation (SD). The carbon in the ‘outer-rhizosphere’ samples was measured in three samples randomly taken from each of the eight pot replicates (see Figure 1 for sampling regime). Relative changes compared to the control are also reported. Significant changes determined by t-tests are highlighted in bold.

| | inner-rhizosphere | | | outer-rhizosphere | | |
|---|---|---|---|---|---|---|
| total C | control | 112 | 1852 | control | 112 | 1852 |
| AV | 1.10 | **1.26** | **1.28** | 1.09 | **1.17** | 1.11 |
| SD | 0.08 | 0.14 | 0.21 | 0.06 | 0.07 | 0.07 |
| % control | | 15.1 | 16.8 | | 8.2 | 2.0 |

SI Table 4: C and N contents and C/N ratio of three soil fractions in control and two isolate treatments. Mean, standard deviation (SD), % difference to control and ANOVA results shown.

| | | carbon | | | nitrogen | | | C/N | | | C/N ratio of C and N increase vs control | | |
|---|---|---|---|---|---|---|---|---|---|---|---|---|---|
| | | control | 1852 | 112 | control | 1852 | 112 | control | 1852 | 112 | control | 1852 | 112 |
| POM | mean | 0.16 | 0.17 | 0.17 | 0.003 | 0.004 | 0.006 | 53 | 42 | 27 | N/A | 12 | 4.4 |
| | SD | 0.07 | 0.09 | 0.06 | 0.002 | 0.001 | 0.002 | 18 | 20 | 1.6 | | | |
| | % control | 0.0 | 7.0 | 8.4 | 0.0 | 28.0 | 93.0 | 0.00 | -21.2 | -48.3 | | | |
| | ANOVA | | | | a | ab | b | a | ab | b | | | |
| AggC | mean | 0.40 | 0.43 | 0.49 | 0.020 | 0.023 | 0.027 | 20 | 19 | 18 | N/A | 9.4 | 12 |
| | SD | 0.06 | 0.07 | 0.05 | 0.004 | 0.005 | 0.004 | 2.00 | 2.74 | 2.19 | | | |
| | % control | 0 | 5.8 | 21.3 | 0.0 | 12.2 | 33.8 | 0.00 | -4.8 | -9.6 | | | |
| | ANOVA | a | ab | b | a | ab | b | | | | | | |
| MAOM | AV mean | 0.46 | 0.51 | 0.50 | 0.034 | 0.039 | 0.039 | 14 | 13 | 13 | N/A | 8.5 | 7.7 |
| | SD | 0.03 | 0.02 | 0.03 | 0.002 | 0.002 | 0.003 | 0.93 | 0.66 | 0.45 | | | |
| | % control | 0.00 | 9.8 | 8.0 | 0.0 | 15.8 | 14.2 | 0.00 | -5.2 | -5.6 | | | |
| | ANOVA | a | b | ab | a | b | b | | | | | | |
